# Rot or not? Uncovering the spatial patterns and drivers of Norway spruce root rot with harvester data

**DOI:** 10.1101/2025.02.07.637055

**Authors:** Susanne Suvanto, Juha Heikkinen, Eero Holmström, Juha Honkaniemi, Tuula Piri, Jarkko Hantula, Tapio Räsänen, Kirsi Riekki, Juha-Antti Sorsa, Harri Hytönen, Henna Höglund, Tuomas Rajala, Aleksi Lehtonen, Mikko Peltoniemi

## Abstract

Root rot is a major problem for forestry, leading to reduced timber quality, growth losses, and increased disturbance risks. Harvester data provides a promising source of information for improving the knowledge on the root rot distribution. Here, we used harvester data (1) to map the risk of spruce root rot in southern and central Finland, and (2) to understand the drivers of the spatial patterns in rot occurrence. First, we built a statistical model predicting the percentage of stems affected by root rot on stand-level. To train the model, we used an extensive set of harvester data, containing 10,402 clear-cut forest stands, where the presence of root rot was recorded for each cut tree using an algorithm based on bucking patterns (i.e., cutting of the stem into different log assortments) recorded by the harvester. The model consisted of two parts, a fixed component describing the effects of different drivers of root rot, and a spatial random component describing the spatial patterns not explained by the fixed part of the model. The fixed part included forest and site attributes, landscape characteristics and proxies of forest-use legacies. The model was then used to map root rot risk, by predicting the probability of root rot occurrence using spatial data sets of the variables in the fixed part of the model, and the known rot status of locations in the data set for the random part of the model. Finally, the map was tested with an independent validation data, verifying its ability to identify the high-risk areas. Proxies of forest-use legacies, tree size and site fertility were found to drive the percentage of rot-affected stems in stands. The results quantify the root rot risk in Finland in higher detail than before and demonstrate the large potential of harvester data in informing about the risk of root rot in boreal forests.

## Introduction

Root rot disease caused by *Heterobasidion* species is a serious problem in coniferous forests throughout the temperate and boreal regions of the Northern Hemisphere (Garbelotto and Gonthier, 2013). In Finland alone, the economic losses from *Heterobasidion* root rot on Norway spruce are estimated to be more than 50 million euros per year, consisting of wood value losses due to decay and the profit losses due to growing less valuable species in the next rotation period (Hantula et al., 2023). In addition, the disease considerably reduces tree growth (Hellgren and Stenlid, 1995; Oliva et al., 2010), making the true economic losses even greater. Furthermore, root rot increases the vulnerability of trees to other damage agents, such as windthrows and bark beetles (Asiegbu et al., 2005; Garbelotto and Gonthier, 2013; Honkaniemi et al., 2017; Wahlman et al., 2025). The warming climate is expected to increase the reproduction and spread of root rot in boreal conditions (Müller et al., 2015, 2014).

In southern Finland, nearly 80% of root rot in Norway spruce (*Picea abies* L.) is caused by fungi from the *Heterobasidion* genus (Piri et al., 1990). The primary species responsible is *H. parviporum* Niemelä & Korhonen, which is a serious cause of decay in spruces and larches, but not a problem for Scots pine. In addition, in spruce trees similar decay as *H. parviporum* may be caused by *H. annosum* (Fr.) Bref. s.s., which has a broader host range, including several broad-leaved and coniferous species, also Scots pine (Korhonen, 1978; Piri et al., 1990). Trees can be infected by *Heterobasidion* via two routes. The primary infection of a tree is caused by airborne basidiospores landing on freshly exposed wood surfaces, such as new stumps. Then, once the fungus is established, it may cause secondary infections by spreading to new trees by vegetative growth of the mycelium through root contacts (Garbelotto and Gonthier, 2013). This secondary infection route can also infect the next tree generation, as the fungus can stay alive and infectious in the stumps decades after the felling (Asiegbu et al., 2005; Piri, 1996). Along with *Heterobasidion* spp., several other fungi cause root and butt rot in Norway spruce, with *Armillaria* species and *Stereum sanguinolentum* being the most common of these (Hallaksela, 1984; Piri et al., 1990).

The presence of root rot in a forest has an impact on the forest management. As *Heterobasidion* root rot can spread to the next tree generation from infected stumps, the tree species selection of the next rotation is a key strategy for controlling the disease (Ahtikoski et al., 2024; Aza et al., 2022; Möykkynen and Pukkala, 2009). Regenerating an infested spruce site with resistant tree species and removing naturally established spruce regeneration can help eradicate the disease (Piri and Korhonen, 2001). The ability of root rot to spread to the new tree generation is also a challenge for continuous cover forestry, as the new spruce regeneration can become infected from diseased overstory trees through root contacts (Piri and Korhonen, 2001; Piri and Valkonen, 2013). Accurate information on the distribution of the *Heterobasidion* root rot is thus needed to support the forest management decisions in high-risk areas in finding the best management approach and species composition.

In Finland, *Heterobasidion* root rot is found mainly in the southern and central parts of the country, especially in coastal areas (Mattila and Nuutinen, 2007; Tamminen, 1985). In northern parts of the country its distribution is limited, and rot is mainly caused by other pathogens. The spread of *Heterobasidion* spp. in the north is most likely limited by loggings historically being conducted during the wintertime, effectively preventing the spread of the species, and also by low soil pH (Müller et al., 2018). On stand level, the incidence of root rot is linked to forest age and size of the trees (Garbelotto and Gonthier, 2013; Piri et al., 1990; Räty et al., 2021). Disease incidence has been observed to be higher on fertile and sandy soils and soils with low organic matter content (Garbelotto and Gonthier, 2013; Mattila and Nuutinen, 2007), whereas the rate of infection has been observed to be lower in peat soils compared to mineral soils (Piri and Vainio, 2024; Redfern et al., 2010). Mixed forests have been hypothesized to have a lower risk of root rot due to fewer root contacts between spruce trees. This was supported by results from Finnish forests, where amounts of damage by *Heterobasidion* root rot were slightly lower in mixed stands compared to pure spruce stands, although other factors were found to be more important for the incidence of rot (Piri et al., 1990). Human activities, including former forest management practices and use, have also contributed to the current distribution of root rot disease. Creation of fresh stumps in summer cuttings creates an effective route for spores to enter new trees, thus creating favourable conditions for the spread of the fungus (Honkaniemi et al., in prep.; Piri et al., 1990). Land use history also plays a role, as rot damage has been found to be generally more severe on former agricultural lands and pastures compared to forest soils (Garbelotto and Gonthier, 2013), although in Finland Piri et al. (1990) did not find a significant relationship between indicators of past land use and the incidence of rot.

Current information on the spatial distribution of spruce root rot in Finland is mainly based on observations from the national forest inventory (NFI) and from field surveys specifically carried out to detect root rot (Mattila and Nuutinen, 2007; Müller et al., 2018; Piri et al., 1990; Piri and Vainio, 2024; Tamminen, 1985). With the NFI data, the challenge is the uncertainty of the observations, as the rot is challenging to detect in living trees (Mattila and Nuutinen, 2007; Tamminen, 1985). On the other hand, while the root rot focused field campaigns targeting recently cut forests can identify rot cases more accurately and produce more detailed information, they are time-consuming and expensive, thus limiting the total sample size that can be attained. More recently, new methods have been tested for identifying rot-affected trees, including remote sensing (Allen et al., 2022; Pitkänen et al., 2021; Räty et al., 2021) and the use of harvester data (Holmström et al., 2024b; Lara et al., 2024; Räty et al., 2021, 2023). Since root rot is best detected immediately after a tree is cut, data collected with a harvester provides a promising approach for collecting information on root rot. Routinely collected rot information with harvesters could enable high density of observations, covering large areas, with the potential of updates when more data accumulates. Yet, the potential of this data source is still largely unexplored.

Here, we analysed data from cut-to-length harvesters, where the rot-impacted stems are identified based on cut patterns, (1) to map the probability of root rot occurrence in southern and central Finland, and (2) to better understand the drivers of the spatial patterns in root rot occurrence, using proxies of past management and forest use legacies, site properties and stand and landscape characteristics. The harvester data set used in the study contained tree-level data from 10,402 forest stands.

## Methods

### Root rot detection algorithm

The root rot detection algorithm automatically identifies stems affected by root rot based on the bucking information (i.e., records from cutting of the stem into different log assortments) recorded by the harvester. For each processed stem, the bucking data produced by the harvester contains information about the bucked logs, including the lengths and top diameter of the log, volume and the log assortment. While the harvester has in-built automation that gives a suggestion about the cutting points for the stem, the machine operator can also deviate from this suggestion. This is typically done when there are defects or quality-issues with the stem.

The algorithm detects the stems affected by root rot based on the log assortment and the location of the log on the stem. It identifies cases where no sawlog has been cut from the first two meters from the butt-end of a sawlog-sized stem, but at least one sawlog has been produced from the upper parts of the stem. This type of bucking pattern is typical in cases where the lower part of the stem is affected by root rot and thus does not qualify as a sawlog. The rot-detection algorithm was developed by Metsäteho and is described in more detail in Holmström et al. (2024b, under review).

### Harvester data and processing

The harvester data set consists of data from clear-cut forest stands in southern and central Finland from 2017 to 2021. The selection of data from a larger database was conducted using weighting to balance the geographical distribution of observations and to increase the representation of spruce-dominated forests (spruce accounted for the highest share of harvested volume in 93% of stands in the data).

Each sawlog-sized spruce stem in the data was accompanied with rot information of either containing rot (1) or being free of rot (0), based on the detection with the root rot algorithm described above (section ‘Root rot detection algorithm’). The logging records were delineated to stands based on the locations of the logged stems with the method of Riekki and Malinen (2022), and stands were only included if they contained at least 100 sawlog-sized spruce stems (i.e., stems to which the rot-detection algorithm was applied to). Stands were excluded from the analysis if any of the predictor variables used in the model was not available (see section ‘Predictor variables’ for details of the variables). For each stand, the percentage of rot-affected stems was then calculated as the share of sawlog-sized spruce stems that contained rot. In total, the final data set consisted of 10,402 forest stands, containing in total 11,774,673 harvested stems, 5,098,578 of which were sawlog-sized spruce stems.

### Predictor variables

The variables tested for predicting root rot contained attributes of the forest stand and site, variables describing the landscape within 1 km radius, and proxies related to the past forest use and management (Table 1). The tested predictors related to the forest stand were calculated from the harvester data and included the average diameter at breast-height of all harvested trees (DBH_mean) and the percentage of spruce from the total harvested volume of the stand. The site variables contained site fertility class extracted from MS-NFI forest resource maps (Mäkisara et al., 2022) and soil type (mineral vs organic) extracted from the topographic data base (National Land Survey of Finland, 2022). The landscape variables included the share of non-forest area and the share of water area from the landscape, calculated from the Corine Land Cover data (Finnish Environment Institute SYKE, 2018) and the share of spruce dominated forests from the forest area in the landscape, based on geospatial data sets of the Finnish Forest Centre (https://www.metsakeskus.fi/fi/avoin-metsa-ja-luontotieto/aineistot-paikkatieto-ohjelmille/paikkatietoaineistot).

**Table 1.**
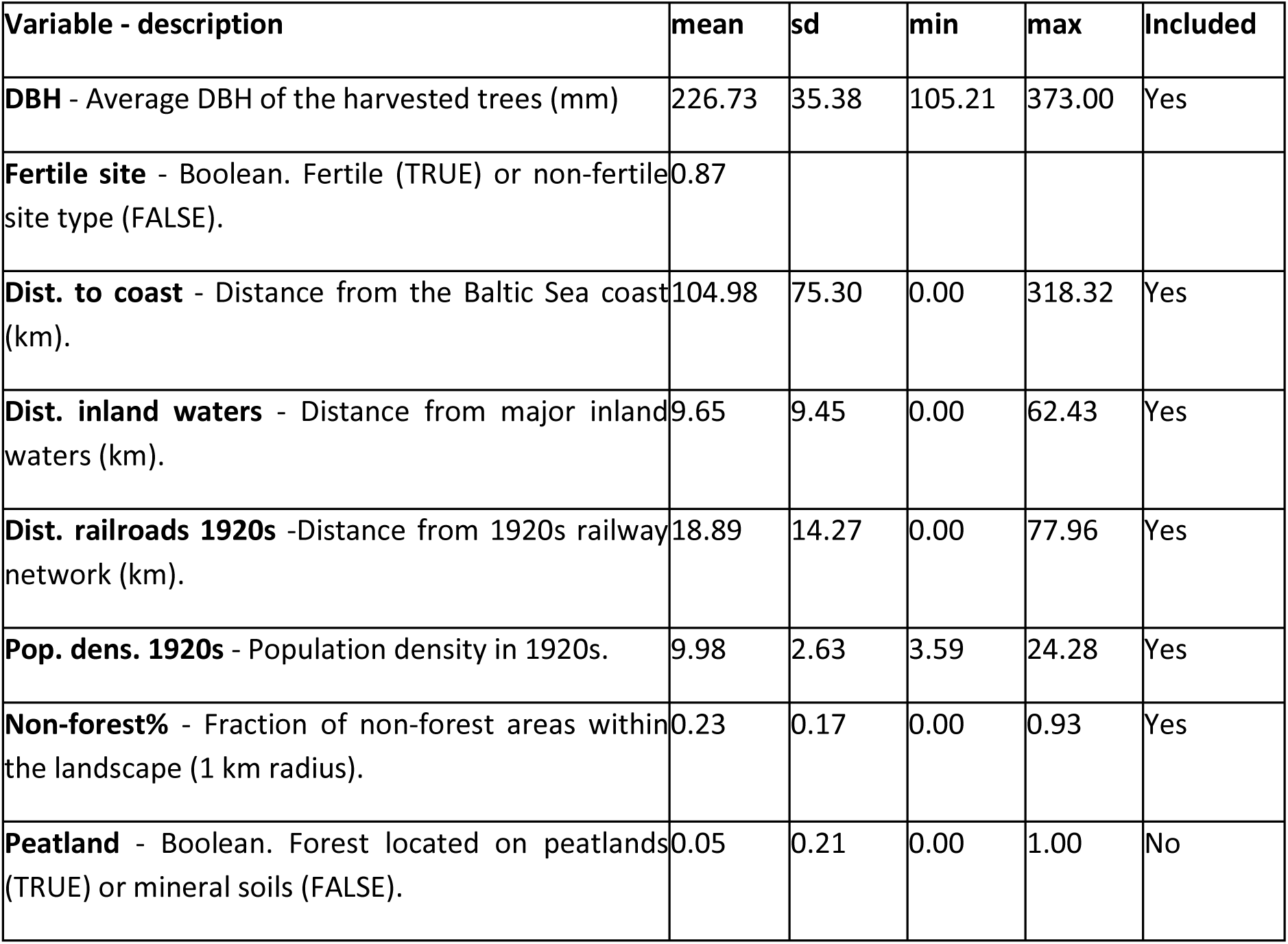
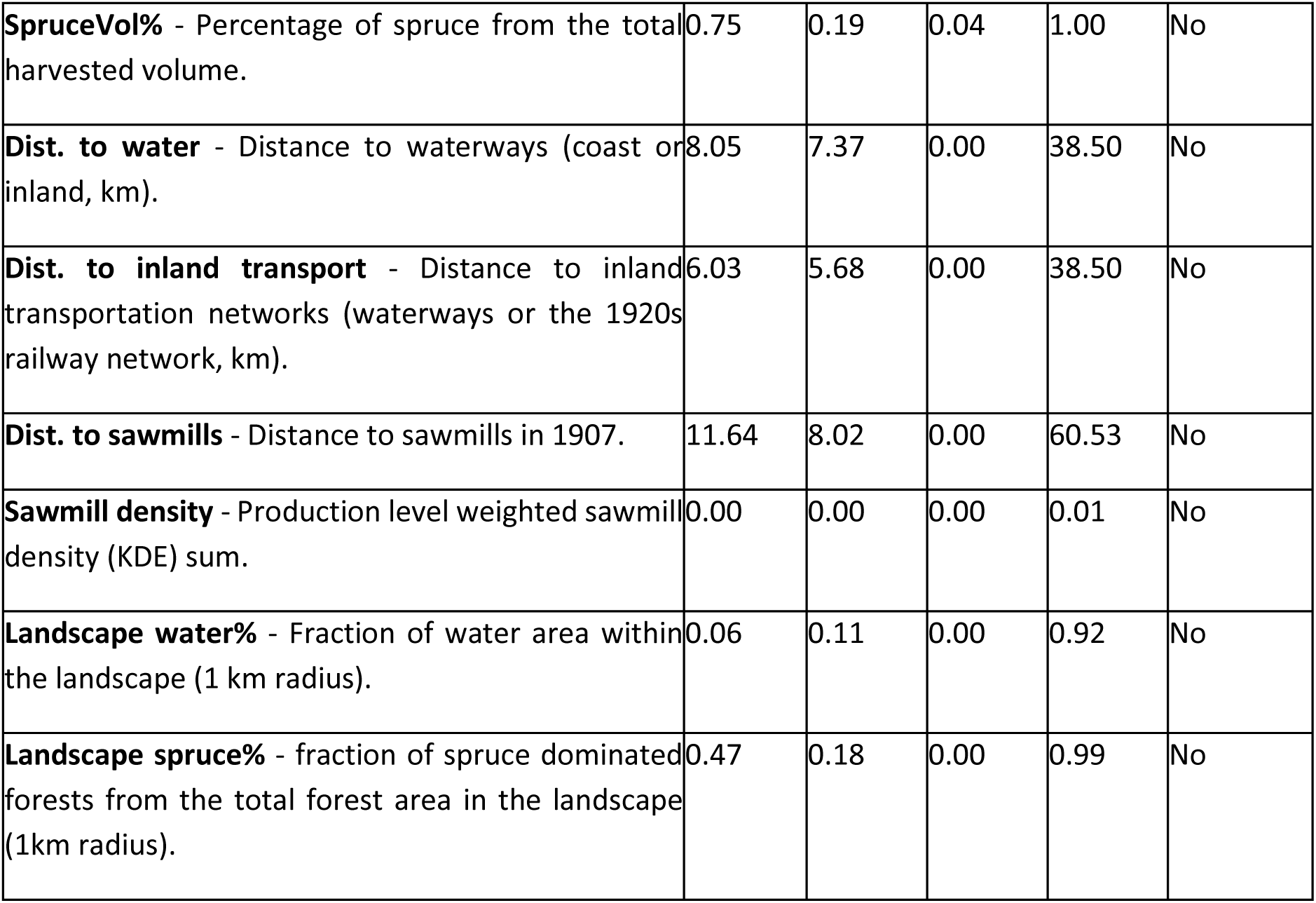
Descriptions and statistics for the predictor variables tested in the model selection, and information on whether or not they were included in the final model. For the boolean variables the reported mean shows the fraction of TRUE observations.

| Variable - description | mean | sd | min | max | Included |
| --- | --- | --- | --- | --- | --- |
| **DBH** - Average DBH of the harvested trees (mm) | 226.73 | 35.38 | 105.21 | 373.00 | Yes |
| **Fertile site** - Boolean. Fertile (TRUE) or non-fertile site type (FALSE). | 0.87 |  |  |  |  |
| **Dist. to coast** - Distance from the Baltic Sea coast (km). | 104.98 | 75.30 | 0.00 | 318.32 | Yes |
| **Dist. inland waters** - Distance from major inland waters (km). | 9.65 | 9.45 | 0.00 | 62.43 | Yes |
| **Dist. railroads 1920s** -Distance from 1920s railway network (km). | 18.89 | 14.27 | 0.00 | 77.96 | Yes |
| **Pop. dens. 1920s** - Population density in 1920s. | 9.98 | 2.63 | 3.59 | 24.28 | Yes |
| **Non-forest%** - Fraction of non-forest areas within the landscape (1 km radius). | 0.23 | 0.17 | 0.00 | 0.93 | Yes |
| **Peatland** - Boolean. Forest located on peatlands (TRUE) or mineral soils (FALSE). | 0.05 | 0.21 | 0.00 | 1.00 | No |
| **SpruceVol%** - Percentage of spruce from the total harvested volume. | 0.75 | 0.19 | 0.04 | 1.00 | No |
| **Dist. to water** - Distance to waterways (coast or inland, km). | 8.05 | 7.37 | 0.00 | 38.50 | No |
| **Dist. to inland transport** - Distance to inland transportation networks (waterways or the 1920s railway network, km). | 6.03 | 5.68 | 0.00 | 38.50 | No |
| **Dist. to sawmills** - Distance to sawmills in 1907. | 11.64 | 8.02 | 0.00 | 60.53 | No |
| **Sawmill density** - Production level weighted sawmill density (KDE) sum. | 0.00 | 0.00 | 0.00 | 0.01 | No |
| **Landscape water%** - Fraction of water area within the landscape (1 km radius). | 0.06 | 0.11 | 0.00 | 0.92 | No |
| **Landscape spruce%** - fraction of spruce dominated forests from the total forest area in the landscape (1km radius). | 0.47 | 0.18 | 0.00 | 0.99 | No |

Proxies for past forest use legacies included historical population density in the 1920s (Wittig, 1928), distance to the railway network in the 1920s (Wittig, 1928) and the inland waterways (i.e., streams with a catchment area of at least 1000 km^2^ or connected lakes; Finnish Environment Institute SYKE, 2021), and proximity to the saw mills operational in 1907 (Alfthan, 1911). In addition, combinations of these variables were also tested, including the distance to waterways (coast or inland) and the distance to inland transportation network (railways or inland waters). The influence of proximity of historical sawmills was explored with two variables: distance to the closest sawmill and the sawmill density, which was calculated as a kernel density estimate with the contribution of each sawmill weighted by their production value. The data from Wittig (1928) was obtained from the digitalised version by Aakala et al. (2023) and data from Alftan (1911) digitalized by Honkaniemi et al. (in prep.). The population density data was provided as point location data, representing a given number of inhabitants. This was processed into a raster following the approach of Aakala et al. (2013) by first aggregating the point data into a 10 km grid and then calculating inverse distance weighing (IDW) interpolation with the *spatstat* package in R (*spatstat.explore* version 3.2-6; Baddeley et al., 2015). Details of the variables can be found in Table 1.

### Statistical analysis

The statistical modelling approach consisted of two steps: (1) fitting a fixed-effects linear regression model to predict the percentage of rot-affected stems, and (2) modelling the spatial structure of the root rot that was not explained by the fixed-effects model as random-effects by fitting a variogram to the residuals of the fixed-effects model. This set-up was chosen to be able to both model the relationship of root rot to specific drivers, but also to account for and utilize the strong spatial autocorrelation pattern of the root rot risk, which could not be fully explained by the predictors of the fixed effects model.

In the fixed-effects model, we applied a log-transformation to the response variable (the percentage of rot-affected stems), and then fitted a linear regression model with independent variables that potentially relate to rot occurrence based on previous literature. All covariates were scaled and centered before the model fitting. Model selection was carried out by comparing alternative model compositions with Akaike’s Information Criteria (AIC) and choosing the variable combination that resulted in lowest AIC values and where all coefficient estimates statistically significantly different from 0 (with p < 0.01). In addition, we considered the plausibility of the effect direction from the point of view of previous understanding of root rot drivers. Different variable combinations were tested iteratively by adding variables to the previously identified best variable combination. For continuous variables with non-negative values, a log-transformation of the variable was also tested and compared with AIC. We calculated the variation inflation factor (VIF) and only allowed variable combinations in the model where VIF was lower than 2 for all predictor variables. After the model selection, the form of the equation for the final fixed-effects model was:

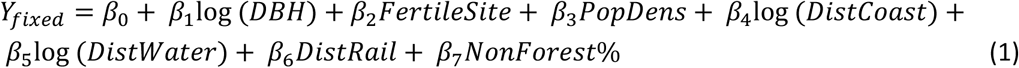

where Y_fixed_ is the percentage of rot-affected sawlog-sized spruce stems in the forest stand (logaritmically transformed). Log(DBH), FertileSite, PopDens, DistCoast, DistWater, DistRail and NonForest% are the scaled and centered predictor variables (see full descriptions in Table 1, in case of log-transformations, the scaling was done after the transformation), and β_0_ to β_7_ are the corresponding coefficient estimates. Please find further information on the reasons for exclusion of different variables in the supplementary material (Table S1).

The random-effects model was then applied to the residuals of the final fixed-effects model, by, first, fitting an empirical variogram with 500-meter bin-width and a 15-kilometre cut-off distance. These parameters were chosen to fit the spatial structure observed in the data. Then, a spherical variogram model was fitted to the empirical variogram. These steps were conducted with R package *gstat* (version 2.1; Pebesma, 2004) using functions *variogram* and *fit.variogram*. With the resulting spatial random-effects model, it was possible to attain both a location-based prediction (Pred_kriging_) and variance (Var_kriging_) for the residuals of the fixed-effect model. Thus, by combining the prediction of the fixed and random effects models, we get a final prediction for the percentage of rot-affected stems, and the variance component of the random-effects model also provides an uncertainty estimate for this prediction.

### Mapping the root rot risk

The rot risk map shows the probability of a forest stand having rot in more than 10% of the sawlog-sized spruce stems, and it is calculated by utilizing the model predictions and the variance component of the random effects model. To map the risk of root rot spatially across the study area, we combined the fixed and random effects models, spatial data sets on the fixed-effect model predictors, and the locations of the harvester data observations. The mapping was done at a 16 m x 16 m spatial resolution to be consistent with the forest resource maps for Finland (Mäkisara et al., 2022).

First, we compiled spatial data describing the predictor variables included in the final fixed-effects model and computed a model prediction for each grid cell. Stem diameter (DBH) was fixed to the average value in the data set (227.4 mm), as the effect of diameter on rot risk is not indicative of whether or not the root rot occurs on the site, but rather whether the rot has reached high enough in the stem to be detectable. Distance to the Baltic Sea, the inland waterways and the historical railway network were calculated by first rasterizing the original vector data to the 16-meter resolution grid and then calculating raster distance (proximity) for this rasterised layer. The processing was done with gdal (GDAL/OGR contributors, 2022) and QGIS (QGIS.org, 2023). Population density was processed from the point data provided by Aakala et al. (2024) into a raster as described earlier (section ‘Predictor variables’). The resolution of all raster layers was resampled to match the final 16-meter resolution of the risk map.

Second, the locations and residuals for the data points in the harvester data set were used together with the spatial random-effects model to compute the random effect prediction (Pred_kriging_) and variance (Var_kriging_) for each grid cell. The full prediction of the percentage of rot-affected stems was then calculated as:

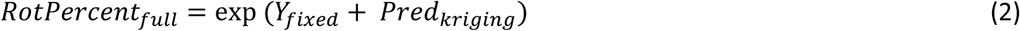

where *RotPercent_full_* is the final prediction of the percent of rot-affected stems in the stand, *Y_fixed_* is the prediction of the fixed-effects model and *Pred_kriging_* is the prediction of the spatial random-effects component.

The final map, representing probability for more than 10% of sawlog-sized spruce stems to have rot (“rot risk map” from here on), was obtained by calculating the value of the cumulative distribution function (CDF) at log(10) of a normal distribution, when the mean is defined as the *RotPercent_full_* and variance as the *Var_kriging_* of each cell location. The value of the CDF then represents the probability of the rot percent being equal to or below 10%, and the rot risk thus is defined as

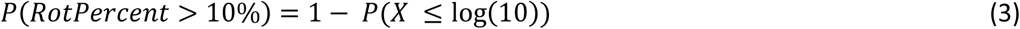

where *P(X ≤ log(10))* is the value of the CDF at log(10).

### Processing to control for privacy and uncertainty

Due to privacy concerns related to the root rot data, we processed the data to make sure that the data point locations could not be derived from the resulting map. Patterns that could lead to identification of data point locations resulted from the random effects part of the map, because the variation of the random-effects (Var_kriging_) would always be lower in the proximity of the data point locations, and in cases with high residuals (i.e., large differences between the fixed-effects model prediction and the observed rot percentage) the random-effect prediction (Pred_kriging_) would also differ distinctively in the proximity of the data point. Therefore, we used a Gaussian kernel (with parameter sigma set to 4000 m) to smooth the Pred_kriging_ raster before using it in the map calculation. For the random-effects variance (Var_kriging_) raster, we excluded the values within a 1.9 km radius of the data points (to cover the range of the fitted variogram model) and replaced these values with the average of the remaining neighbourhood cells.

Map values were only presented for areas with reasonable density of harvester-data observations, by excluding the grid cells where less than 5 observations were present in the data set within a 15 km radius.

### Validation

The rot risk map was validated with independent test data that was not included in the model development. This data set was processed and filtered with the same criteria as the original data set, by excluding any stands with fewer than 100 sawlog-sized spruce stems. The filtered test data consisted of 533 stands, of which 25 were excluded as they were not included in the final risk map area after the observation density mask had been applied. The final test data therefore contained 508 stands, consisting of in total 230,916 sawlog-sized spruce stems (i.e., stems for which the rot detection algorithm was applied). The test data had approximately the same geographical distribution as the model training data, and the distributions of average stem diameter of harvested trees and the share of spruce of total harvested volume were also comparable to the training data.

The validation was carried out by calculating the risk map values for the locations of the test data and comparing these with the observed rot information in the data points, i.e. whether more than 10% of the stems in the stands had rot. When calculating the map values for the locations, we used the observed average diameters in the test data (DBH) and not the fixed stem diameter value as in calculating the final rot risk map.

## Results

### Factors explaining root rot in stands

From the variables related to the forest and site characteristics, the percentage of rot-affected stems was most related to the average DBH of the harvested trees, with higher rot probability in forest stands with higher DBH, and site type, with higher rot probability in the more fertile sites. Forest stands located in landscapes with a high percentage of non-forest area were also found to have more rot. Root rot was found to be strongly related to variables representing past forest-use legacies. The variables in the final fixed-effects model include the 1920s population density and the distance to the Baltic Sea coast, distance to the inland waterways and distance to the 1920s railway network, which are all proxies for connectivity to historical timber transportation networks (Fig. 1A, Table 2). More information about the results for variables tested but not included in the final fixed effects model is included in the Supplementary material (Table S1).

**Figure 1.**
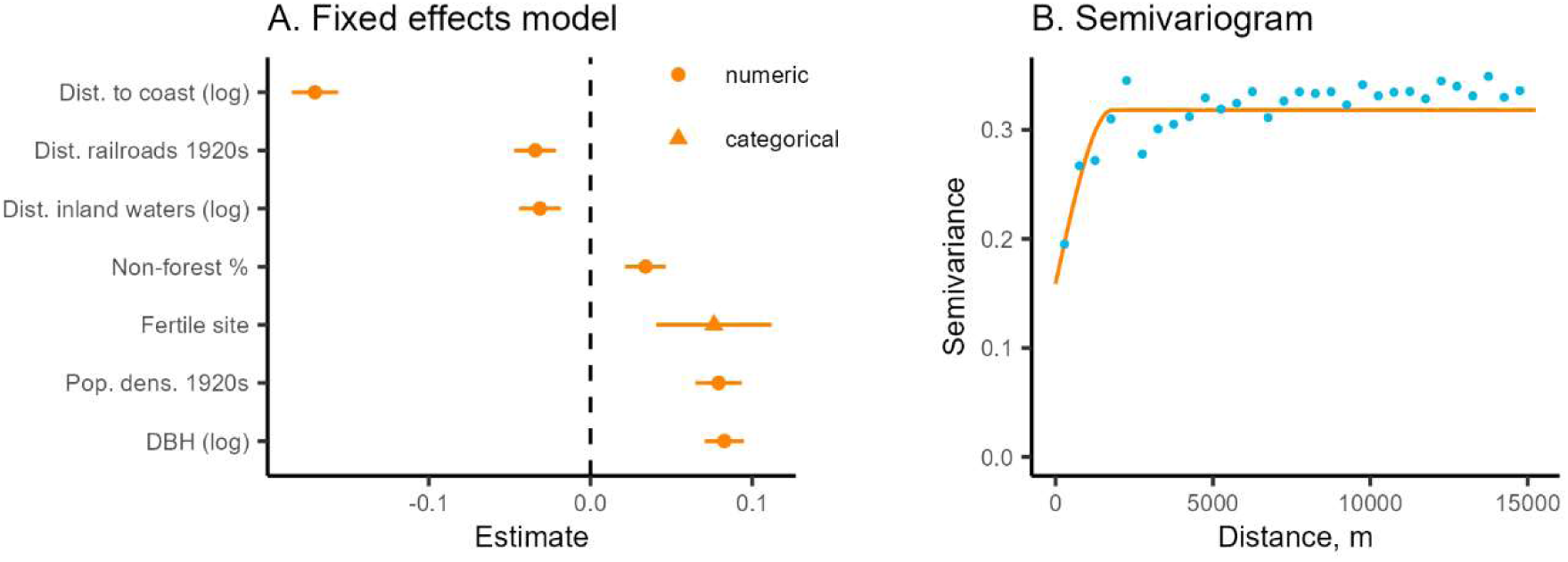
Coefficient estimates and their 95% percent confidence intervals for the fixed effects model (A), and the variogram of the (spatial) random effects model (B).

**Table 2.** The coefficient estimates (Est.) and their p-values (p) for selected alternative models including variable for peatland soils (peat vs. mineral soil, variable “Peatland”) as a predictor (columns Peat 1-4), and for the final fixed-effects model used for creating the rot risk map (Final Model). All models are fit to the same training data.

|  | Peat 1 |  | Peat 2 |  | Peat 3 |  | Peat 4 |  | Final Model |  |
| --- | --- | --- | --- | --- | --- | --- | --- | --- | --- | --- |
|  | Est. | p | Est. | p | Est. | p | Est. | p | Est. | p |
| (Intercept) | 2.407 | <0.001 | 2.403 | <0.001 | 2.400 | <0.001 | 2.320 | <0.001 | 2.336 | <0.001 |
| DBH (log) |  |  | 0.080 | <0.001 | 0.108 | <0.001 | 0.086 | <0.001 | 0.083 | <0.001 |
| Fertile site |  |  |  |  |  |  | 0.088 | <0.001 | 0.076 | <0.001 |
| Pop. dens. 1920s |  |  |  |  |  |  | 0.080 | <0.001 | 0.079 | <0.001 |
| Dist. coast (log) |  |  |  |  | -0.213 | <0.001 | -0.173 | <0.001 | -0.170 | <0.001 |
| Dist. inland waters (log) |  |  |  |  |  |  | -0.032 | <0.001 | -0.031 | <0.001 |
| Dist. Railroads 1920s |  |  |  |  |  |  | -0.033 | <0.001 | -0.034 | <0.001 |
| Non-forest% |  |  |  |  |  |  | 0.037 | <0.001 | 0.034 | <0.001 |
| Peatland | -0.091 | 0.003 | -0.024 | 0.432 | 0.058 | 0.045 | 0.121 | <0.001 |  |  |

The percentage of rot-affected was not found to be lower at peatlands compared to mineral soils when other predictor variables were also included in the model (Table 2). When the impact of the peatland variable on rot percent was assessed alone, peatlands did have significantly lower rot percentages compared to mineral soils (Table 2, model Peat 1). However, if average diameter of the harvested trees (DBH) was also included in the model, the effect of the peatland variable became non-significant (model Peat 2). Moreover, when both DBH and distance to the Baltic Sea coast were included in the model with the peatland variable, the coefficient estimate for peatland flipped to positive, i.e. indicating more rot on peatlands compared to mineral soils (model Peat 3). This effect became even stronger when the peatland variable was added to the final fixed-effects model containing more variables (model Peat 4). However, as the impact of the peatland variable in the model was opposite to what we expected based on previous research, it was left out from the final model used for the rot-risk map (see more details in the Discussion section).

The spatial random-effects model component, fitted to the residuals of the fixed-effects model, showed strongest spatial correlation structure within approximately a 1.8 km distance (Fig. 1B, variogram range 1802 m, with nugget 0.16 and sill 0.32).

### Spatial patterns of the root rot risk

The rot-risk map, presenting the probability for observing rot in > 10% of the sawlog-sized spruce stems, shows clear spatial variation (Fig. 2A). Highest probabilities for are found close to the southern coast of the Baltic Sea, around and especially east from Helsinki. Other hotspots are found on the south-western coast (around Turku), along the coast of Gulf of Bothnia (Ostrobothnia, around Vaasa) and in the Tampere region more inland. In these hotspot regions, the probabilities were high, mostly falling into the two highest risk categories (80-90% or >90% probability). The fine-scale variation in the rot-risk values is low (Fig. 2B), with the map mainly showing gradients of changes in the risk across larger distances, whereas the local differences are lower and mainly arise from the site fertility variable.

**Figure 2.**
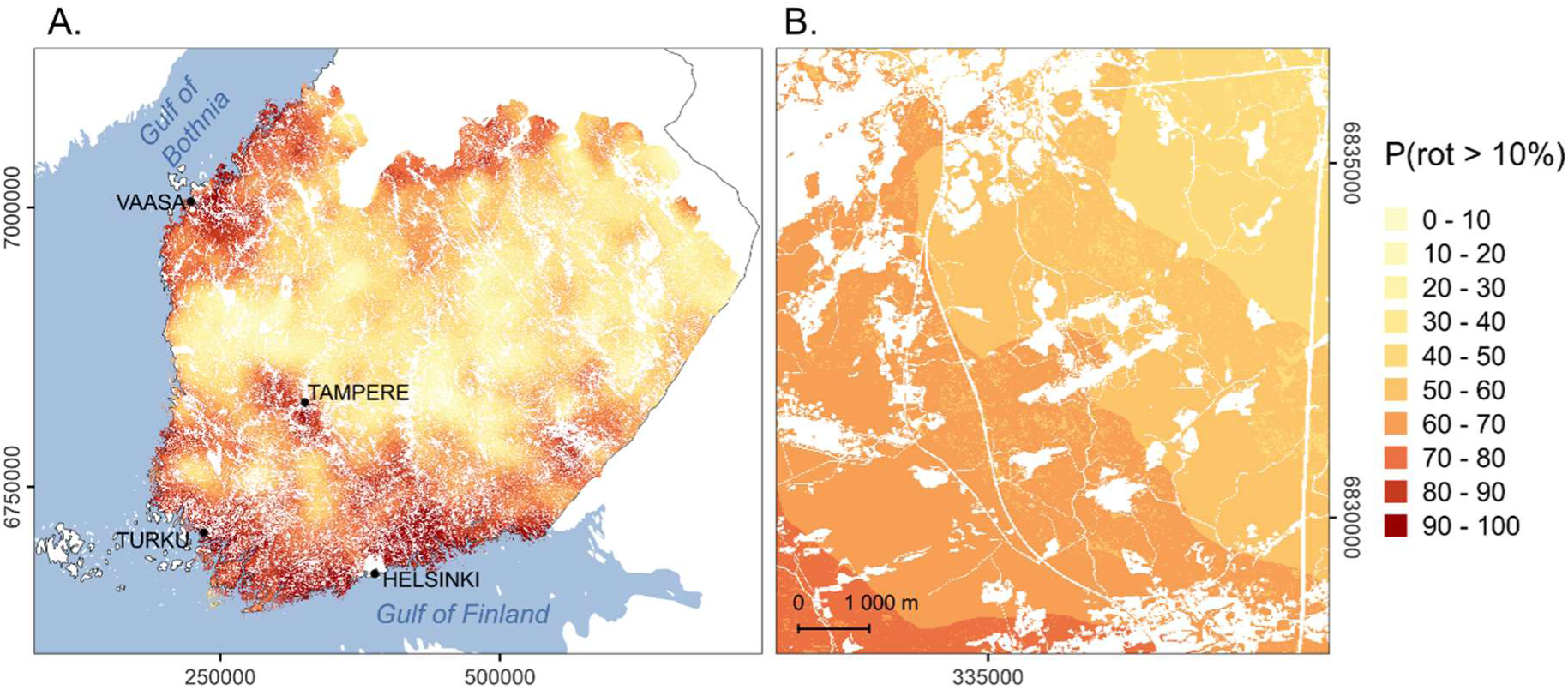
The risk map, showing the probability of more than 10% of sawlog-sized spruce stems to be rot-affected (A) and a fine-scale detail of the map (B). White colour on the map represents non-forest areas.

### Validation

The validation of the risk values (i.e., probability of RotPercent exceeding 10%, from Eq. 3) suggested that the map was able to identify high-risk areas, especially when the fixed and random effects model components were combined. The AUC value comparing the predicted probabilities and the observed rot cases (>10% of stems) was 0.73, whereas the AUC value for the fixed-effects model predictions only was lower, 0.69 (Fig. 3A). The frequencies of the observed rot cases corresponded well with the rot risk classes on the map in the test data locations when both fixed and random effects model components were used, whereas the fixed-only predictions were not as well aligned with the test data (Fig. 3B). The validation of the model predictions for the percentage of stems affected by rot (RotPercent, from Eq. 2), combining the fixed and spatial random effects components, resulted in a RMSE of 7.50% (8.09% for fixed-model only) and R-squared of 0.26 (0.14 for fixed-model only).

**Figure 3.**
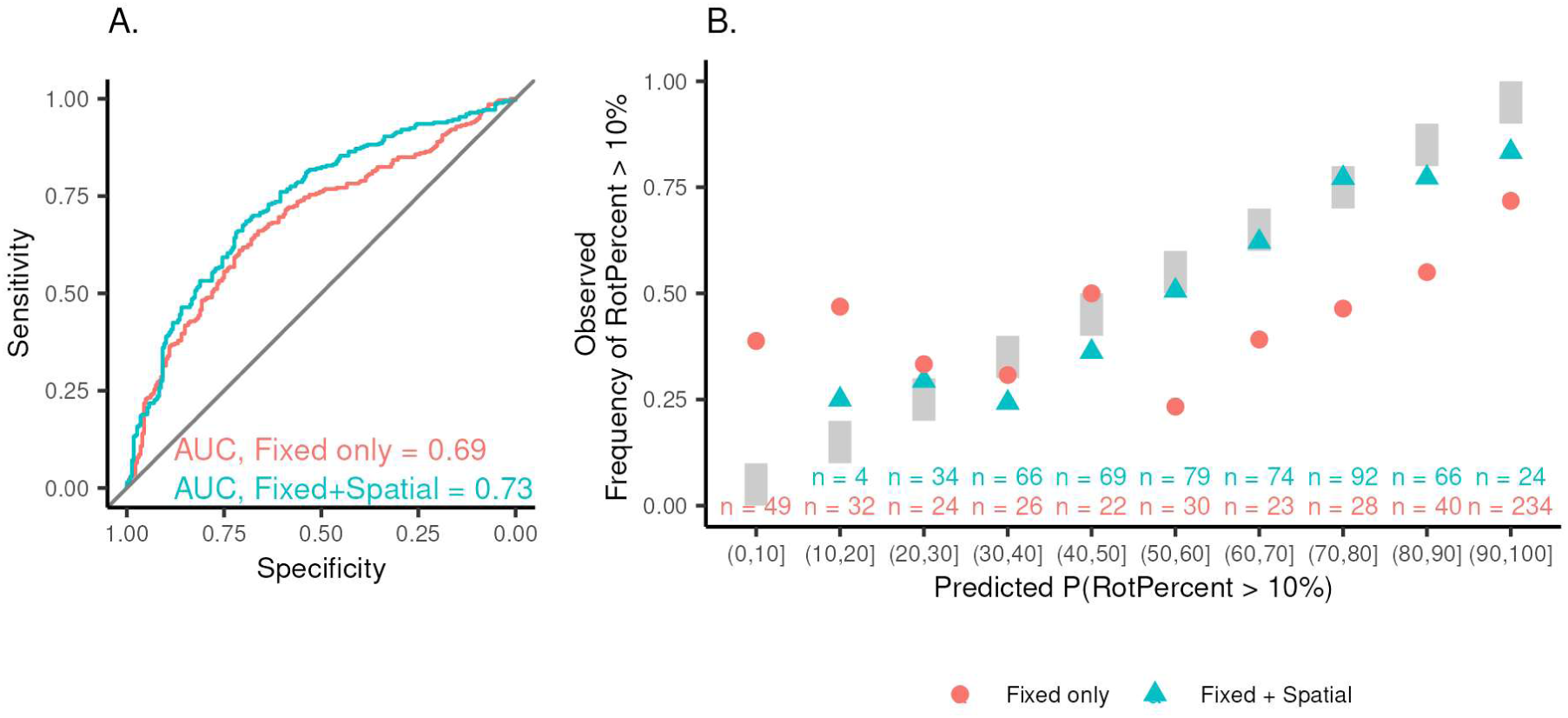
Validation of the map with the external test data showing: (A) the ROC curves and AUC values for the mapped risk values (i.e., P(RotPercent > 10%) and (B) the observed share of rot cases in each rot risk category of the map. Both results are presented for the fixed model only (red color) and the full predictions containing both the fixed and the spatial component (blue color). In subfigure B, the grey blocks illustrate the range of values included in each category, and the black dot the share of rot cases (>10% of stems identified to have rot) from all stands within this class in the test data.

## Discussion

### Mapping the risk of root rot

The use of harvester data to map root rot risk in our work enabled deriving improved information about the distribution and drivers of root rot. While the previous often NFI-based quantifications of large-scale spatial patterns of root rot risk in Norway spruce have showed comparable spatial patterns (Mattila and Nuutinen, 2007; Tamminen, 1985), the use of harvester data and the automated rot detections allows a great increase in the number of tree-level observations to be used. For example, the NFI-based analysis by Mattila and Nuutinen (2007) was based on observations from 8 007 trees measured in the NFI, compared to more than 5 million trees run through the rot-detection algorithm for this work. Futhermore, when the rot is observed during the harvest operation (as the bucking decision based on the visual interpretation of the machine operator), it is less likely to go undetected compared to the observations of root rot in the NFI data where the detection of rot in living trees can be challenging (Mattila and Nuutinen, 2007; Tamminen, 1985). While field-campaigns specifically focusing on mapping root rot on recently cut sites do not have this problem, they are often limited to relatively small sample sizes, due to the costs of the field work. Deriving rot-information from harvester data overcomes both issues.

The validation results showed that the prediction of the exact percentage of rot-affected spruce stems in an individual forest stand is still challenging. Therefore, instead of presenting the predicted percentage of rot on the map, we utilised the variance estimate from the spatial random effects model component and mapped the probability for more than 10% of the spruce stems to have rot. The results were found to agree well with the observed frequencies of forests in the validation data set where the number of rot-affected stems exceeded the 10% threshold (Fig. 3). Thresholding probabilities is a common step in binary classification, and the threshold selection is always case dependent. Here, the 10% threshold was selected based on the consultation of forestry experts and advisors who considered that above this threshold, owners and buyers are typically concerned about root rot. By quantifying the risk as a probability for exceeding 10% of rot-affected stems we were able to quantify the risk with reasonable accuracy, even if the prediction of the exact rot-percentage was not feasible.

The risk map shows clear large-scale variation in the root rot risk across the study area (Fig. 2A). The overall spatial patterns in our results, with high rot-risks along the coast, are well in line with the results from previous studies (Mattila and Nuutinen, 2007; Tamminen, 1985), but provide a more detailed picture of the spatial variation in rot risk. The variation on the map at the finest scale remained low (Fig. 2B) and mainly results from the variation in site type (variable “Fertile site”). The other forest and site-specific variables were fixed on the map (average DBH of harvested trees) or excluded from the final model, whereas other predictors used for the included in the model described broader spatial patterns. Model prediction was highly impacted by the average DBH of harvested trees, which – if included in the map calculation – would lead to higher variation between forest stands in local scales. However, DBH was held constant on the map (fixed to the average of the data), as we expect the higher rot incidence with higher DBH to be caused by the rot having more time to spread in the stem, whereas smaller trees may be infected by even if rot is not yet detected in the stem. Thus, letting the DBH vary across the landscape in the map prediction would tell more about where the rot has already reached the stem, not where the trees are infected. However, for applications where the prediction is needed for a specific tree size , e.g. for predicting areal decay losses of wood, one can apply the fixed-effects model together with the prediction from the spatial random effects model component (see Data availability statement ).

The rot status in near-by stands, incorporated into the map through the spatial random-effects component, was crucial for predicting the risk of root rot. This is even though we purposefully degraded the level of detail presented due to privacy reasons (as described in section ‘Processing to control for privacy and uncertainty’). In spite of degrading the spatial details on the map, the validation still confirms the ability of the map to identify the risk of root rot, in which the random-effects component played a crucial role (Fig. 3). This implies that incorporating the information of the rot status from known locations in the neighbourhood crucially improves the prediction, when compared to using only the predictions of the fixed-effects model. Similar conclusions were drawn by Räty et al. (2021), who found that the error rates for the predictions of root rot in Norway spruce were considerably higher when cross-validation was conducted with spatially distinct clusters.

The current risk map has potential to be improved significantly in the future as more harvester data is accumulated. As harvester data is automatically collected with new cuttings, the data source has great potential to be updated and extended on continuous basis. The increasing number of observations would improve the accuracy of the map, but also reduce the need for degrading the map details due to privacy reasons, as the higher observation density would obscure the distinctive patterns around individual data points. In addition, future development in detection of rot-affected stems during the harvest operation, such as automated image-based detection of rot (Holmström et al., 2024a; Ostovar et al., 2019), could further improve the use of harvester data in the mapping of root rot.

### Drivers of root rot risk

Site fertility and tree size (average DBH) were the two site-specific variables included in the fixed effects model for predicting the percentage of rot-affected stems. The impact of these variables, and the direction of the effect, i.e., more rot on fertile sites and more rot with increasing tree size – are in line with previous research on the topic (Garbelotto and Gonthier, 2013; Mattila and Nuutinen, 2007; Piri et al., 1990). The effect of tree size for rot incidence is very clear and it has been established in several studies with different kinds of approaches, including field research focused on (Piri et al., 1990), NFI-based studies (Mattila and Nuutinen, 2007; Thor et al., 2005), and in research utilizing harvester data (Räty et al., 2021).

The effect of species composition was not statistically significant in our results. In previous research (Piri et al., 1990), mixed forests have had slightly lower damage levels observed, but the relationship between species composition and rot was not particularly strong. Honkaniemi et al. (in prep.) and Thor et al. (2005) found the percentage of spruce in the stand to increase occurrence in the NFI plots in Finland and Sweden, respectively. As noted by Piri et al. (1990), the current species composition may not correspond to the species composition at the time of the infection. Even current monospecific stands may have been developed from previously mixed-species stands, as thinnings typically have favored conifer species while broad-leaved trees have been removed. The disconnection between the observation and infection time forest structure applies also to many other forest attributes that can be correlated with the incidence (except for tree size, as this has a direct relationship to incidence, as the rot has had more time to advance in older and larger trees).

Surprisingly, our results did not suggest a higher root rot risk for mineral soils compared to peatlands, even though *Heterobasidion* caused is considered to be less common in peatlands (Mattila and Nuutinen, 2007; Piri and Vainio, 2024; Redfern et al., 2010). While our data showed significantly higher percentages of rot-affected stems in mineral soils, this effect disappeared when tree size was considered, and the effect further changed to the opposite direction when other variables were included in the model (Table 2). The cause for this observed pattern is not clear, but there are a few perspectives to consider. First, our data does not allow the identification of the fungal species responsible for the detected rot. Compared to mineral soils, in peatlands is less frequently caused by *Heterobasidion* sp. and more often by other wood-decaying fungi, such as *Armillaria* sp. (Piri and Vainio, 2024). This suggests that also in our results the rot on peatland is likely more often caused by other species than *Heterobasidion* sp.. On the other hand, it is possible for the automated rot-detection algorithm to erroneously identify cases of other stem damage or quality defects as rot. The validation of the rot-detection algorithm, reported by Holmström et al. (2024b), showed that while well-developed rot in stems was easily identified correctly by the machine operators, smaller-scale rot cases were sometimes mixed with other quality defects or stem damage. Therefore, if other stem quality defects were more common on peatlands compared to mineral soils, this could lead to higher frequency of wrongly detected rot and cause the observed pattern in our results. However, there is no clear evidence of higher frequency of stem form defects in peatland spruce forests compared to mineral soils (Rikala, 2003; Stöd et al., 2002). Finally, the forest use history between forests on peatlands and mineral soils is different, with the active forestry becoming increasingly common on peatlands during the latter half of the 20th century, as demonstrated by the drastic decrease in undrained peatlands within this time (Korhonen et al., 2024). As active forest use history increases the probability of root rot infection and the spread of the fungus is slow, it is possible that the infections initiated after mid-20th century have now developed to a stage where they can be observed in final fellings. Additional uncertainties in our results are caused by not having field-verified information about the soil type at the sites, but we had to instead use the peatland data in the national-level topographic database, which may not always accurately represent the local conditions at the sites. Peatland forests were also underrepresented in the data base, with only 5% of the data being located on peatlands, whereas in total approximately one quarter of forests in Finland are peatland forests (Korhonen et al., 2024). For these reasons, we have decided to exclude the peatland variable from the final model used for creating the rot risk map, as it was not clear if the results provide enough evidence for representing higher rot risk on peatlands. Regardless of this, the results raise a clear need to investigate the situation of peatlands forests in more detail in the future.

Many of the variables selected into the final model were proxies of past forest-use legacies. This is in line with previous research linking the spread of the *Heterobasidion* root rot to past forest and land use (Garbelotto and Gonthier, 2013; Piri et al., 1990), and results relating NFI-based rot observations to proxies of past forest-use (Honkaniemi et al., in prep.), and supports the assumption that past forest-use legacies have played a major role in shaping the present patterns of incidence in Finland. The contributions of past forest use are challenging to interpret in detail, because proxies of past forest use are typically correlated with other variables, in particularly climate. Historically the human population and activities have concentrated into the southern parts of the country and the coastal areas, which are also the areas with higher average temperatures.

The limited availability of information about the past forest use also makes it difficult to study their impacts on the distribution of rot. For example, here we found the distance to the Baltic Sea coast to perform the best in the statistical model, compared to more detailed historical variables derived from the locations of historical sawmills, which were also heavily concentrated on the coastal areas (more details of excluded variables in the supplementary material S1). As the spatial patterns of forest use have varied in time and by the type of forest use (Tasanen, 2004), it may be that the distance to the coast serves as a better overall proxy for the past forest use activity, whereas the variables derived from the historical sawmill data describe only one forest use type at one point in time. Yet, the interpretation of the distance to the coast is less clear. Indeed, this pattern of higher rot incidence on the coast has been detected before, and it has been attributed, for example, to climatic effects (Mattila and Nuutinen, 2007) and to higher wind load on coastal areas, potentially leading to more windbreaks that providing entry points for the fungi that cause root rot (Tamminen, 1985). However, considering the spatial patterns of forest wind damage in Finland as observed from the NFI data (Suvanto et al., 2019) the latter hypothesis seems unlikely. In our results the plausibility of the impact of past forest use as the key driver is supported by the significance of other human-related proxy variables (historical population density; distance to inland transportation routes, railways and inland waterways) that have fewer alternative explanations.

### Implications for forest management

The presence of in a forest impacts the optimal forest management decisions to be made, both in terms of species selection when the forest is regenerated (Ahtikoski et al., 2024; Aza et al., 2022; Holmström et al., under review) and in terms of the type and scheduling of the planned harvest operations (Aza et al., 2021; Möykkynen and Pukkala, 2009; Piri and Vainio, 2024). In particular, practicing continuous-cover forestry in forest stands infected by is challenging, as infection spreads from the previous tree generation to the new tree generation, and frequent harvests provide opportunities for new primary infections (Nevalainen, 2017; Piri and Korhonen, 2001; Piri and Valkonen, 2013). The results of this work provide an updated and detailed spatial estimation of the risk of across Southern and Central Finland that can support forest owners and managers in taking the root rot risk into account in forest management. Therefore, we believe it can become an important tool for forest owners and buyers, as material steering the education, dissemination, and forestry guidance activities and campaigns to high-risk areas.

The created data set can also help in taking into consideration when assessing the future role of different forest management approaches in meeting different goals set to forests. For example, continuous-cover forestry (CCF) has been seen as a major tool for balancing biodiversity goals, wood production and other ecosystem services in forest management (Blattert et al., 2023; Eyvindson et al., 2021; Peura et al., 2018), but these analyses have so far not considered root rot risks, even if CCF likely should be avoided in rot-risk areas (Piri and Korhonen, 2001; Piri and Valkonen, 2013). Accounting for the impacts of on the value of the harvested wood and the growth of the trees could give an improved assessment on the potential of CCF management at high-risk areas.

## Conclusions

Our results produced a quantification of root rot risk in southern and central Finland with improved level of detail, and demonstrated the potential of harvester data and automated rot-detection for mapping of risk. Use of harvester data enables the collection of large amounts of observations from cut trees during harvesting with rot-information. This reduces the probability of rot being undetected at individual stands, and gives a more realistic understanding of the general rot status of regions, as it is challenging to identify it from living trees (e.g. in national forest inventories). Furthermore, accumulating harvester data from new harvest events presents a possibility for updating the map and improving its quality as an increasing number of observations becomes available. With the presented modelling framework, the new harvester data could easily be used to update the risk map through the spatial random-effects component even without necessarily needing to re-fit the models. Considering the current harvesting rates in Finland and the fact that modern harvesters are already producing the data needed for the analysis, it would be possible to create a considerably improved map after accumulating harvester data for a few years.

The study provided important insights into the factors behind the observed root rot situation. While the results for, for example, stand attributes and site fertility were in line with the earlier understanding, we surprisingly found peatland forests to have an increased risk of root rot, when other tree size and location were controlled for. This raises a clear need for further investigations of the root rot situation of peatland forests in the future. Our study also showed evidence that the past forest use has a profound influence on the root rot incidence, impacting more broadly also the current forest damage regime, where root rot plays a significant role through increasing the probability of wind damage and bark beetle incidences. These results emphasize the long timescales over which the current distribution of root rot has been formed and the difficulty of understanding the root rot status based on the current forest characteristics alone. On the other hand, the fixed model was only able to explain a limited amount of the spatial distribution of root rot risk, whereas the spatial random component was found to be crucial for the predictions. This means that observations about the rot-status of near-by forests are needed for a reliable assessment of the rot risk.

## Supplementary material

Supplementary material is available attached at the end of this preprint file, containing correlation matrices between the response and predictor variables in the models (Fig. S1) and additional information on the model selection process (Table S1).

## Acknowledgements

The authors wish to acknowledge CSC – IT Center for Science, Finland, for computational resources.

## Data availability statement

The resulting data products from this work, including the root rot risk map and the raster data layers used for its calculation, will be available openly in Zenodo at the time of the publication of the paper with DOI 10.5281/zenodo.14825296. The original harvester data underlying the work is used with an agreement from third party data owners and cannot therefore be openly shared by the authors.

## Supplementary material

**Figure S1.**
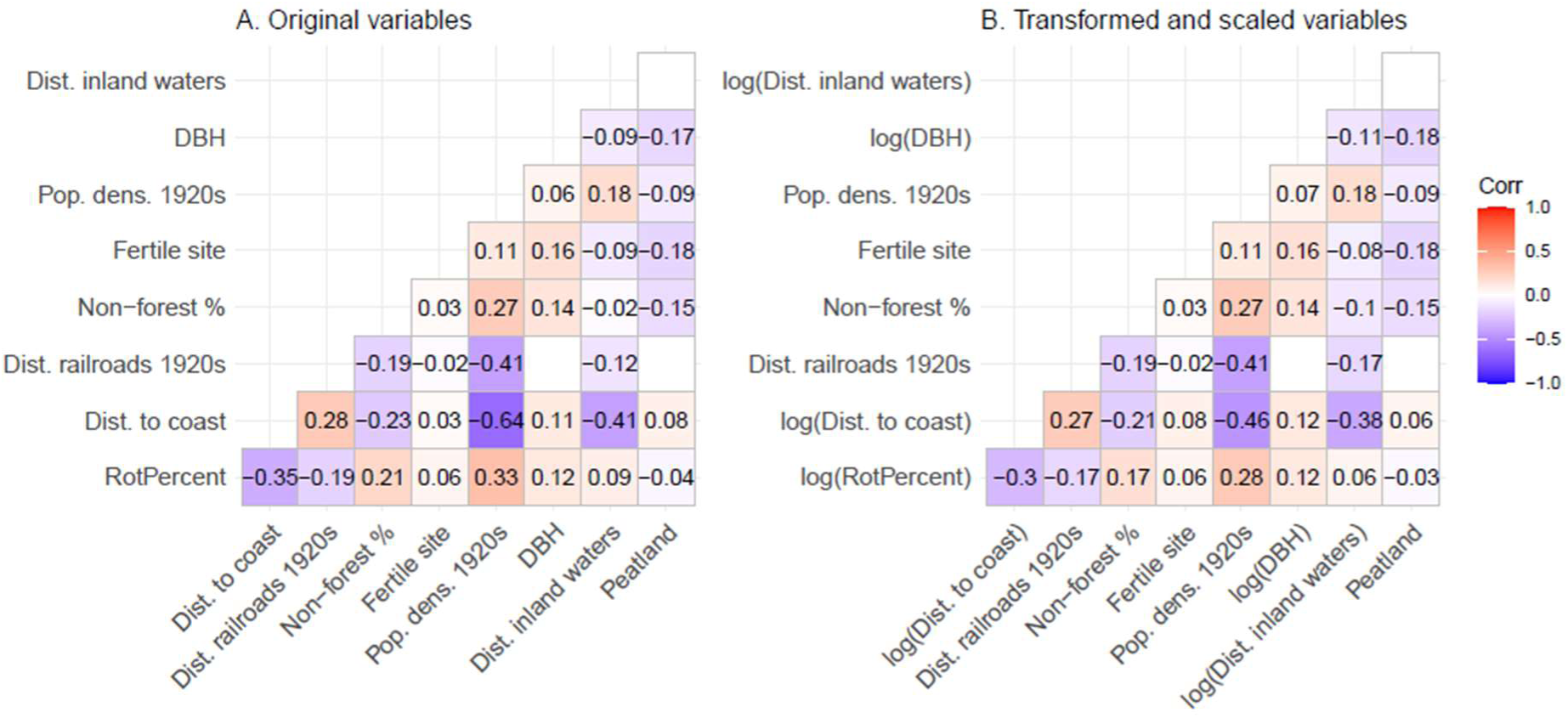
Correlation matrices for root rot percent (RotPercent), the variables included in the final model, and the Peatland variable explored in the alternative models. Subfigure A shows the correlations of variables without transformations, and B shows the correlations with the transformations included in the final model. Numeric values are only shown for statistically significant correlations (p < 0.001).

**Table S1.**
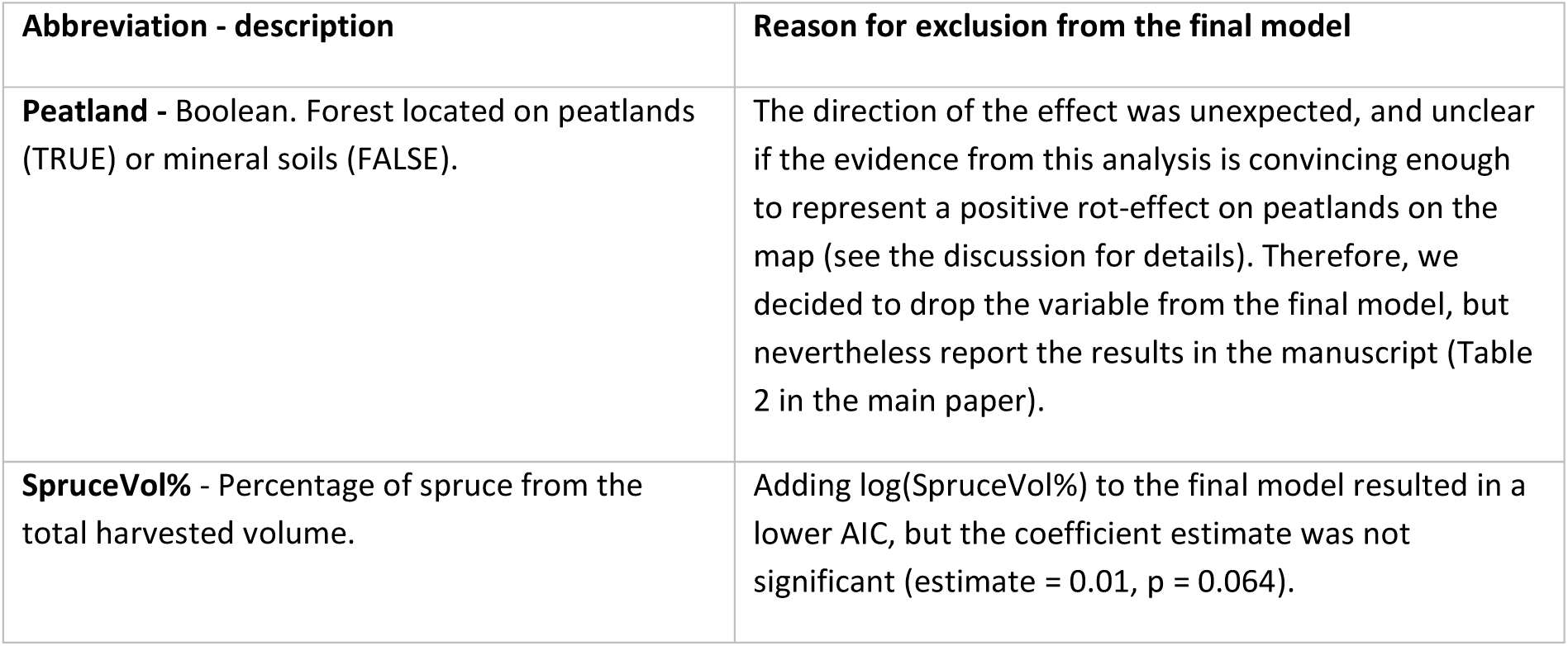
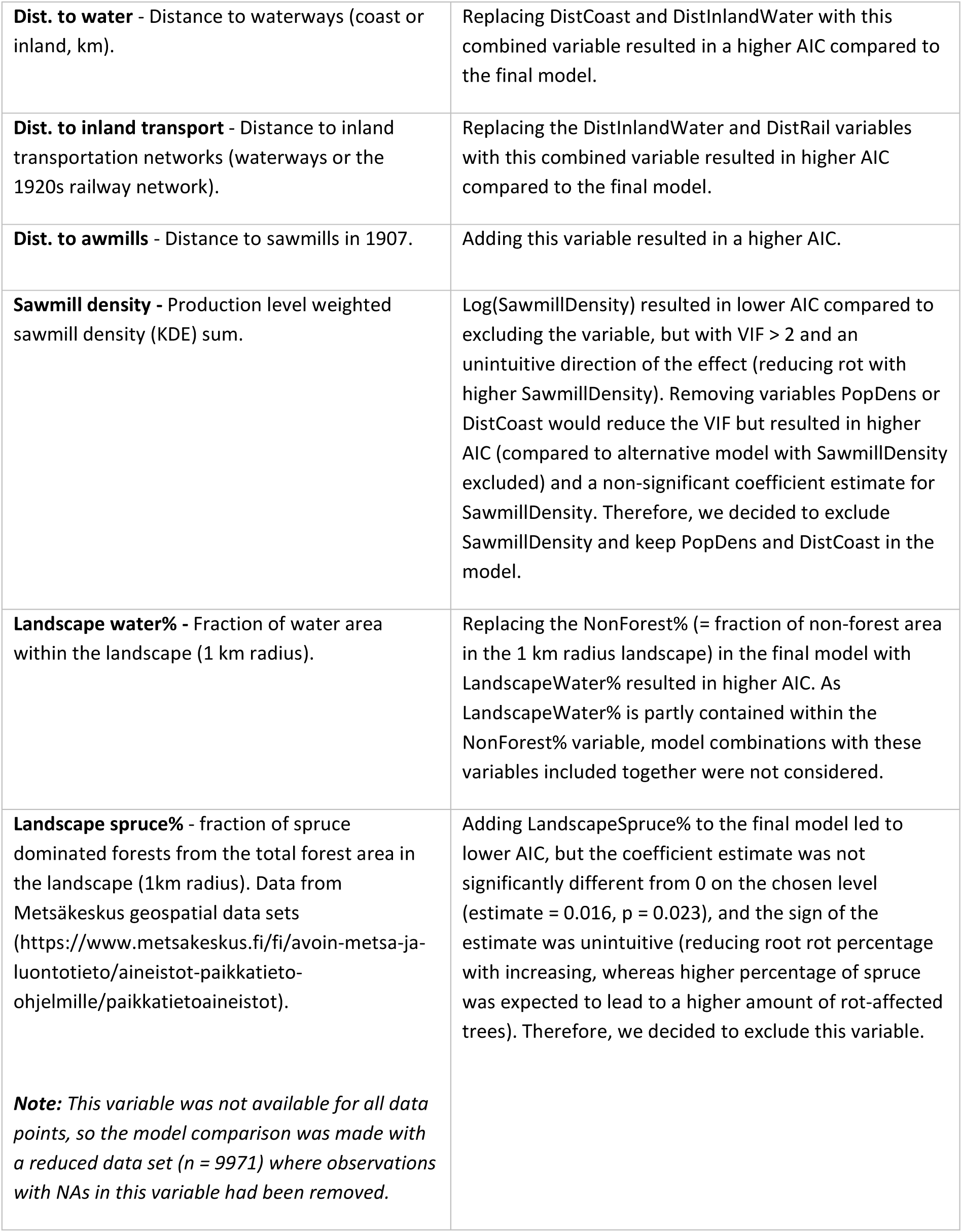
Details on the reasons for excluding variables from the final model. The model selection was made based on comparison of AIC values, statistical significance of coefficient estimates (p < 0.01) and the plausibility of the relationships (e.g., if the effect in the model was supported by theory).

